# TFEB and MCOLN1 are important for *Coxiella burnetii* egress via lysosomal exocytosis

**DOI:** 10.64898/2026.05.07.723496

**Authors:** Sven Rinkel, Jan Schulze-Luehrmann, Fiona Weber, Elisabeth Liebler-Tenorio, Anja Lührmann

**Author notes:** **Correspondence:** Prof. Dr. Anja Lührmann, Mikrobiologisches Institut – Klinische Mikrobiologie, Immunologie und Hygiene Universitätsklinikum Erlangen, Friedrich-Alexander-Universität Erlangen-Nürnberg, Wasserturmstraße 3/5, 91054 Erlangen, Germany.

## Abstract

*Coxiella burnetii* is a Gram-negative, obligate intracellular pathogen and the causative agent of the zoonotic disease Q fever. Resident alveolar macrophages are the first target cells, but *C. burnetii* spreads to other cell types. While we have information about *C. burnetii* uptake and the establishment of the replication-competent phagolysosomal-like *C. burnetii*-containing vacuole (CCV), it is not well studied how *C. burnetii* exits its host cell.

Here, we show that an infection with *C. burnetii* also triggers the activation of TFEB, a master regulator of autophagy and lysosomal development. The activation occurs in a time-dependent manner and depends on the size of the CCV. Importantly, TFEB activation during *C. burnetii* infection depend on MCOLN1, which channels Ca^2+^ across the lysosomal membrane into the cytosol. Knock-down of MCOLN1 resulted in reduced TFEB activation and smaller CCVs, while MCOLN1 activation boosted bacterial egress. Indeed, peripheral CCVs are positive for LAMP1/2 and release bacteria, without inducing host cell death. Importantly, LAMP1/2 and *C. burnetii* were stainable in non-permeabilized cells at sites of bacterial release, demonstrating fusion of the lysosome with the plasma membrane. Importantly, while replication of *C. burnetii* is not inhibited in cells lacking LAMP1/2, egress is impaired.

Taken together, our data indicates that with increasing CCV size, TFEB is activated by the release of Ca^2+^ from lysosomes via the MCOLN1 channel, which in turn enables further CCV development and damage of the CCV membrane. This triggers lysosomal exocytosis and egress of *C. burnetii* without cell death induction.

## INTRODUCTION

*Coxiella burnetii* is an obligate intracellular zoonotic pathogen, which infects a variety of different vertebrates and non-vertebrates, such as mammals, birds, reptiles and arthropods (Angelakis *et al*., 2010). Ruminants, like cattle, sheep and goats, are considered its primary reservoir (Maurin *et al*., 1999, Pouquet *et al*., 2020). Infected ruminants might develop coxiellosis, which has a quite diverse manifestation, ranging from being asymptomatic to abortion or reproductive disorders (Agerholm, 2013, Bauer *et al*., 2023). The infected animals shed the bacteria via feces, milk and birthing products. Aerosolized bacteria from these sources are the main cause for human disease (Delsing *et al*., 2011). However, consumption of raw milk products might also cause infection, but less efficiently (Sobotta *et al*., 2025, Miller *et al*., 2021). *C. burnetii* causes Q fever in humans, which can be either acute or chronic (Maurin *et al*., 1999). In addition, Q fever can result in the development of the Q fever fatigue syndrome (QFS), which reduces the patient’s quality of life for months or even years (Morroy *et al*., 2016). The first target cells of *C. burnetii* are alveolar macrophages, but during the course of infection, other cell types also become infected, including endothelial cells, epithelial cells, fibroblasts and trophoblasts (Maurin *et al*., 1999). In these cells, *C. burnetii* establishes a phagolysosomal compartment called the *C. burnetii*-containing vacuole (CCV) (Howe *et al*., 2010, Pechstein *et al*., 2018). The acidic conditions within the CCV lumen activate the type IVB secretion system (T4BSS), which injects effector proteins into the host cell cytoplasm to modulate host cell function (Beare *et al*., 2011, Carey *et al*., 2011). Around 150 effector proteins have been identified (Lührmann *et al*., 2017). For several effector proteins a function in establishing and maintaining the CCV has been assigned (Bauer *et al*., 2023). This is in line with the observation that the T4BSS is essential for the maturation of the CCV into a very spacious vacuole permissive for bacterial replication (Carey *et al*., 2011, Howe *et al*., 2010). However, which host cell factors are involved in the establishment of the spacious CCV is less understood. It was demonstrated that the interaction with the autophagy pathway (Thomas *et al*., 2020), the secretory pathway (Campoy *et al*., 2011) and recycling endosomes (Hall *et al*., 2024) are important for CCV expansion. In addition, the transcription factor EB (TFEB) seems to be involved in the biogenesis of the spacious CCV (Padmanabhan *et al*., 2020). TFEB belongs to the microphthalmia-associated transcription factor (MiTF) family of basic helix-loop-helix (bHLH) transcription factors (Kim *et al*., 2021). They recognize the Coordinated Lysosomal Expression and Regulation (CLEAR) element present within 200 base pairs of the transcriptional start site in many lysosomal (Sardiello *et al*., 2009) and autophagy-related genes (Settembre *et al*., 2013). Hence, TFEB is a *bona fide* master regulator of lysosomal biogenesis and autophagy induction (Raben *et al*., 2016). There are controversial reports about the function of TFEB during *C. burnetii* infection. Initially, it was shown that the infection with *C. burnetii* induces TFEB activation in a T4BSS-dependent manner. Silencing of TFEB resulted in reduced CCV sizes (Padmanabhan *et al*., 2020), indicating a role of TFEB for CCV expansion. Similar results were obtained in another study, where the authors demonstrated that knockouts of TFEB and TFE3, which also belongs to the MiTF family and functions as a *bona fide* master regulator of lysosomal biogenesis, in macrophages led to smaller CCVs (Larson *et al*., 2019). However, in the latter study, the *C. burnetii*-mediated activation of TFEB was independent of the T4BSS. Interestingly, TFEB/TFE3 knock-out cells supported increased bacterial replication, indicating that TFEB might have a different function with regard to CCV maturation and bacterial replication (Larson *et al*., 2019). In contrast, a recent study showed that *C. burnetii* actively blocks TFEB activation in a T4BSS-dependent manner. TFEB-deficient cells were characterized by increased CCV size and improved bacterial growth (Kilips et al., 2024). From these three studies it is still not clear whether i) TFEB is activated or inhibited by *C. burnetii* infection; ii) TFEB activation results in increased or decreased CCV size; iii) the T4BSS is involved in TFEB activation.

Hence, we addressed the question which role TFEB plays in maturation of the CCV and *vice versa*. In addition, we determined how *C. burnetii* egresses after the completion of its replication cycle within the spacious CCV. We had demonstrated recently, that during later stages of infection, when the spacious CCV has been generated, cells might undergo apoptosis, which allows release of infectious particles and the dissemination of the infection (Schulze-Luehrmann *et al*., 2024). However, egress also occurs in cells lacking a functional intrinsic apoptosis cascade, suggesting additional egress strategies.

## MATERIALS AND METHODS

### Reagents and antibodies

Unless otherwise stated, chemicals were purchased from Merck (Darmstadt, Germany) or Carl Roth (Karlsruhe, Germany). The following inhibitors were used: Torin 1 (Cell Signaling, Leiden, Netherlands) and ML-SA1 (Biomol, Hamburg, Germany). The LAMP1 (1D4B) and LAMP2 (ABL-93) specific primary antibodies were developed by J.T. August and obtained from the Developmental Studies Hybridoma Bank (University of Iowa, Department of Biology, Iowa City, IA, USA). Primary antibodies against TFEB, pTFEB (S122 and S211), mTOR, pmTOR (S2448), pRaptor (S792), PRAS40, pPRAS40 (T246), 4E-BP1, p4E-BP1 (T37/46), PP2A C subunit, Mucolipin-1, Pan-Calcineurin A were supplied by Cell Signaling. Primary antibodies against Neuraminidase-1 and Galectin-3 were from Proteintech (Planegg, Germany), and the antibody against Actin was purchased from Sigma Aldrich (Darmstadt, Germany). Secondary antibodies for immunofluorescence staining conjugated with Alexa Fluor-488 or - 594 as well as HRP-conjugated secondary antibodies for immunoblots were from Dianova (Hamburg, Germany).

### Bacterial strains

*Coxiella burnetii* strain Nile Mile Phase II clone 4 (RSA 439), its Δ*dotA* derivative strain (Schäfer *et al*., 2020) and a recently created AnkG deficient strain - Δ*ankG* (Cordsmeier *et al*., 2022) were employed for infection experiments. The *C. burnetii* wild-type strain expressing GFP (*Tn1832*) and a strain lacking a functional type IV secretion system Δ*dotA* GFP (*Tn514*) for microscopy experiments in fixed samples were generously donated by Dr. Matteo Bonazzi (CNRS, University of Montpellier, France) (Martinez *et al*., 2016, Martinez *et al*., 2014). For time-lapse live cell microscopy, *C. burnetii* expressing mCherry (Schulze-Luehrmann *et al*., 2024) and for ratiometric pH measurements *C. burnetii* expressing IPTG-inducible TagRFP were used (Schulze-Luehrmann *et al*., 2016). Bacterial strains were inoculated at 1×10^6^ ml^−1^ and grown in acidified citrate-cysteine medium (ACCM-2) for five days under microaerophilic conditions at 2.5% O_2_, 5% CO_2_ and 37°C. For ACCM-2 plates, 0.3% agarose was added to the liquid medium (Omsland *et al*., 2009). Bacteria cultured on ACCM-2 plates for CFU counts were incubated for 10 to 14 days under microaerophilic conditions.

### Cell lines and culture conditions

EA.hy926 (human umbilical vein endothelial cells), CRL-2992, LOT 63396642 were purchased from ATCC. HeLa (human cervical adenocarcinoma derived epithelial cells) were kindly provided by Prof. Michael Hensel, University Osnabrück. Germany. Both cell lines were cultivated in Dulbeccós modified Eaglés medium (DMEM, Invitrogen, Darmstadt, Germany), 10% FCS (Biochrom, Darmstadt, Germany) at 37°C and 5% CO_2_. Mouse embryonic fibroblasts (MEFs) flox/flox and lamp1/2^−/-^ were kindly provided by Prof. Paul Saftig (Christian-Albrechts-University, Kiel) (Eskelinen *et al*., 2004) and cultured in DMEM, 5% FCS at 37°C and 5% CO_2_.

HeLa cells stably transfected with pWHE644/655-NLS-AnkG cells were cultivated in DMEM, 10% FCS, 1% Penicillin/Streptomycin (Thermo Fisher Scientific), 0.25 µg/ml puromycin (Sigma Aldrich) and 300 µg/ml G418 (Roth) at 37°C and 5% CO_2_ (Berens *et al*., 2015, Pechstein *et al*., 2020).

### Torin 1 and ML-SA1 treatment

HeLa cells were infected with *C. burnetii* for 6 h. The cells were washed several times with PBS, supplied with fresh medium containing 250 nM Torin 1 (Cell Signaling) or 25 µM ML-SA1 (Biomol) and incubated for 18 h at 37°C, 2.5% O_2_ and 5% CO*2*. CFUs of the supernatant and infected cells were performed as described.

### Immunoblotting

Protein samples for detection of mTOR and phospho-mTOR (S2448) were separated by SDS-PAGE on a NuPAGE Tris-Acetate 3-8% gradient gel (Thermo Fisher Scientific). Transfer was performed by wet transfer onto a 0.45 µM PVDF membrane (Merck). All other proteins were separated by SDS-PAGE on a Bolt Bis-Tris Plus 4-12% gradient gel (Thermo Fisher Scientific) at 90V for 120 min and transferred via wet transfer to a 0.45 µM PVDF membrane (Merck) or to a nitrocellulose membrane (Cytiva, Freiburg, Germany). Proteins were detected using the indicated specific primary and corresponding HRP-conjugated secondary antibodies and visualized by a chemiluminescence detection system (Thermo Fisher Scientific).

### Indirect immunofluorescence and live cell imaging

Transfected or infected cells, seeded in a 24 well plate on coverslips, were fixed with 4% paraformaldehyde (PFA, Alfa Aesar, Ward Hill, MA, USA) in PBS (Biochrom) for 15 min at room temperature in the dark. After permeabilization with ice-cold Methanol for 30 s, which was skipped for the Exocytosis staining in MEFs the free aldehyde groups were quenched by a 30 min incubation with 50 mM NH_4_Cl (Roth) in blocking buffer (PBS/ 5% goat serum; Thermo Fisher Scientific) at room temperature. The coverslips were incubated in blocking buffer for 2 h with the primary antibodies indicated, washed three times with PBS and incubated with the corresponding secondary antibodies in blocking buffer for 30 min. After washing three times with PBS the coverslips were mounted using ProLong Diamond with DAPI (Thermo Fisher Scientific).

If necessary and indicated the F-actin cytoskeleton was stained with phalloidin Alexa Fluor 647 (Thermo Fisher Scientific). In this case the permeabilization after fixation with PFA was done by a 5 min incubation in 0.1 % Triton X-100 in PBS instead of Methanol.

For staining with TFEB antibodies, permeabilization was done using 0.1% Triton X-100 in PBS for 10 min. In all following steps - quenching, first and secondary antibody incubation - 0.2% saponin (Alfa Aesar) was added.

Analysis was performed using the Carl Zeiss LSM 700 Laser Scanning Confocal Microscope, recording data with a 63x/1.4 oil immersion objective lens and Zen 3.0 SR (black) software (Carl Zeiss, Oberkochen, Germany).

For live cell imaging, 2 × 10^5^ EA.hy926 cells. infected for 4 days with *C. burnetii* mCherry, were seeded in a 35 mm glass bottom dish No 1.0 (MatTek, Ashland, MA, USA) in 2 ml DMEM, 10% FCS. Twenty-four hours post-seeding SiR actin (Spirochrome AG, Stein am Rhein, Switzerland) was added at 50 nmol in fresh cell culture media and different regions of the infected cells were visualized every 15 min for 12 h with a Zeiss Spinning Disc Axio Observer Z1 (Carl Zeiss).

### TFEB Fluorescence Ratio Analysis

A semi-automated script was created in ImageJ to calculate the fluorometric ratio of nuclear versus cytoplasmic TFEB localization. For this, the cell area of an individual cell, excluding the CCV, was manually outlined. Cytoplasmic measurement was performed only on pixels without overlapping DAPI signal. The nucleus was outlined and measured separately in a second step. The mean TFEB fluorescence intensity per pixel was measured for both compartments. The ratio of nuclear to cytoplasmic TFEB signal was then calculated to assess differential subcellular localization.

### CCV measurement

Immunostaining with anti-LAMP2, anti-TFEB antibodies and DAPI was performed as described above. Combined tile scan and Z-stack pictures were taken with the LSM700 (Zeiss). The volume was calculated using the formula for a triaxial ellipsoid. Measurements of the radii were performed using the Zen 3.0 SR (black) software (Zeiss).

### Ratiometric pH measurements with LysoSensor Yellow/Blue DND-160

EA.hy926 cells were infected with *C. burnetii* expressing IPTG-inducible TagRFP fluorescent protein at MOI 200 for 5d. Then, 2×10^5^ infected cells were seeded in fresh DMEM 10% FCS containing 1 mM IPTG (Thermo Fisher Scientific) 18 h before imaging in 35 mm glass bottom culture dishes (Mat Tek). After washing with equilibration buffer (5 mM NaCl, 115 mM KCl, 1.2 mM MgSO_4_, 25 mM MES; pH 6.0), the cells were incubated in 2 μM Lysosensor Yellow/Blue DND-160 (Thermo Fisher Scientific) in equilibration buffer, and live cell images were acquired between 1 and 15 min after addition of the dye with an ApoTome and Zen software (Carl Zeiss). Cells were excited with 365 nM, and the emission filters were 445 nM (blue) and 510 nM (yellow). Mean pixel values of fluorescence intensities of regions of interest (10 µm^2^) corresponding to CCVs (red fluorescence) were measured using the Zen software (Carl Zeiss). To calibrate Lysosensor Yellow/Blue DND-160 emission ratios for a range of pH values in situ, cells were incubated before addition of the dual-wavelength fluorophore for 15 min with equilibration buffer of known pH (3.8–7.5) in the presence of 10 μM monensin and 10 μM nigericin (Thermo Fisher Scientific) to bring the cytosol and chromaffin vesicles to the pH of the surrounding media. An independent standard curve was generated on each experimental day in Prism to assign a predicted pH for relevant Y/B values. The pH of the cytosol was compared with the pH values of central CCVs and peripheral CCVs from duplicate groups of 30 individual cells in three independent experiments.

### MCOLN1 siRNA knock-down

MCOLN1 siRNA, non-target siRNA and DharmaFECT 1 were purchased from Dharmacon. The siRNAs were diluted in 1x siRNA buffer and incubated for 5 min. Subsequently, the siRNA solutions were mixed with an equal volume of DMEM medium without serum containing DharmaFECT 1, and incubated at room temperature for 20 min. The same volume of HeLa cells in DMEM with 20% FCS was added to the ntRNA- or siRNA-DharmaFECT 1 mixtures. The cells were seeded in 24-well plates and 24 h post-siRNA transfection, the cells were washed with PBS and fresh media with or without *C*. *burnetii* was added.

### Colony-forming units (CFU) count of egressed bacteria and microscopic analysis from *C. burnetii*-infected MEFs

LAMP1/2 knock-out MEFs and corresponding wild-type MEFs were seeded 18 h before infection in either 24-well plates with coverslips at a density of 3 × 10^4^ ml^−1^ for microscopy or in 12-well plates at a density of 5 × 10^4^ ml^−1^ for CFU counting. They were infected with *C. burnetii* or a *dotA*-deficient strain at a MOI of 200 and cultured for 2 days. After five times washing with PBS, the infected cells were incubated with 200 µg/ml gentamicin in cell culture medium for 4 h at 37°C, 5% CO_2_ to kill extracellular bacteria. After washing with PBS, fresh complete media was added to the cells. The coverslips were stained at day 4 post-infection as described before with or without permeabilization with antibodies against LAMP1, LAMP2, *Coxiella* and Phalloidin. For CFU counts of egressed bacteria the supernatants from the 12-well plates were harvested carefully and the pellets (1 min, 20,087 x g) were resuspended in ACCM- 2. Serial dilutions were pipetted in triplicate on ACCM-2/ 0.3% agarose plates, incubated for 10 d at 37°C, 5% CO_2_ and 2.5% O_2_ before CFUs were counted. For intracellular bacteria enumeration, the remaining cells were washed once with 1 ml distilled water and then lysed by addition of 2 ml distilled water for 30 min at RT, followed by repetitive pipetting, transfer to a 2 ml reaction tube and 30 s sonification. Serial dilutions were plated as described above. The percentage of egressed bacteria were calculated by dividing the CFU of egressed bacteria through the total CFU of egressed and intracellular bacteria, multiplied by one hundred.

### Transmission electron microscopy

Wild-type MEFs were infected with *C. burnetii* at MOI 50 for 8 h. The cells were washed several times and fresh medium was added. Cells were prepared for transmission electron microscopy 7 days post infection. For this, the cells were briefly washed with PBS and fixed with 2.5% glutaraldehyde in cacodylate buffer (0.1 M, pH 7.2) for 6 h at 4°C. The fixative was replaced by cacodylate buffer and MEFs were gently scraped off and transferred to an Eppendorf tube. The cells were centrifuged for 5 min at 1,500 x g and the cell pellet was embedded in 2% agarose and sectioned to 1 mm^3^ cubes. Cubes were post-fixed in 2% osmium tetroxide and embedded in Araldite Cy212. Relevant areas were selected in Toluidine-blue-stained semi-thin sections. Ultrathin sections (85 nm) were stained with uranyl acetate and lead citrate and examined in a transmission electron microscope (TECNAI 12, FEI Germany, Dreieich, Germany) at an acceleration voltage of 80 KV, and findings documented using a digital camera (TEMCAM FX416, TVIPS, Gauting, Germany).

### Statistical analysis

Statistical analysis was conducted with Prism 8 (GraphPad software). Bar graphs depict mean data ± standard deviation from three independent experiments in duplicates. An unpaired Student’s t-test or the Mann-Whitney test was performed to determine significance of the results, and a *p*-value of < 0.05 was considered significant.

## RESULTS

### Endogenous TFEB is activated in a time-dependent manner during *C. burnetii* infection

First, we aimed to determine whether *C. burnetii* infection induces or inhibits TFEB activation. In the aforementioned publications, TFEB knock-down and knock-out cells and overexpression of GFP-tagged TFEB were mainly used to elucidate the function of TFEB during *C. burnetii* infection (Killips *et al*., 2024, Larson *et al*., 2019, Padmanabhan *et al*., 2020). Therefore, we decided to study the activation status of endogenous TFEB in cells infected with *C. burnetii* wild-type, the Δ*dotA* mutant, or the Δ*ankG* mutant. The Δ*dotA* mutant is unable to translocate effector proteins (Beare *et al*., 2011, Carey *et al*., 2011), and was therefore used to determine the role of the T4BSS in modulating TFEB. The Δ*ankG* mutant was shown to be affected in the ability to inhibit host cell apoptosis and to generate a replicative CCV (Cordsmeier *et al*., 2022). It was therefore used as a model of a defective CCV. HeLa cells were either infected or not for 72 hours and the subcellular localization of TFEB was determined by immunofluorescence analysis. As shown in figure 1A, TFEB was mainly localized in the host cell cytoplasm in uninfected cells. Similar results were obtained in cells infected with the Δ*dotA* mutant. In contrast, all cells infected with wild-type *C. burnetii* showed a clear nuclear localization of TFEB. The Δ*ankG* mutant seems to only partially activate TFEB, as demonstrated by reduced nuclear localization of TFEB in infected cells. From these data we concluded, that the infection with *C. burnetii* activates TFEB and that bacteria unable to generate a spacious CCV do not. It has been shown that nuclear translocation of TFEB occurs in a time-dependent manner (Padmanabhan *et al*., 2020). To confirm this observation and to determine at which time point of infection the Δ*ankG* mutants starts to deviate from the wild-type with respect to TFEB activation, immunofluorescence images of HeLa cells infected for 24, 48 and 72 hours were analyzed for the subcellular localization of TFEB. In nearly 50% of the cells infected with the wild-type, TFEB was found nuclear at 24 hours post-infection (Fig. 1B). This percentage increased over the infection time period analyzed to nearly 100%. The Δ*dotA* mutant did not induce nuclear localization of TFEB at any tested time point. In contrast, the Δ*ankG* mutant induced nuclear localization of TFEB in around 30% of the infected cells. This percentage did not significantly increase over time. This data indicates that only wild-type *C. burnetii* induced efficient activation of TFEB in a time-dependent manner.

**Figure 1:**
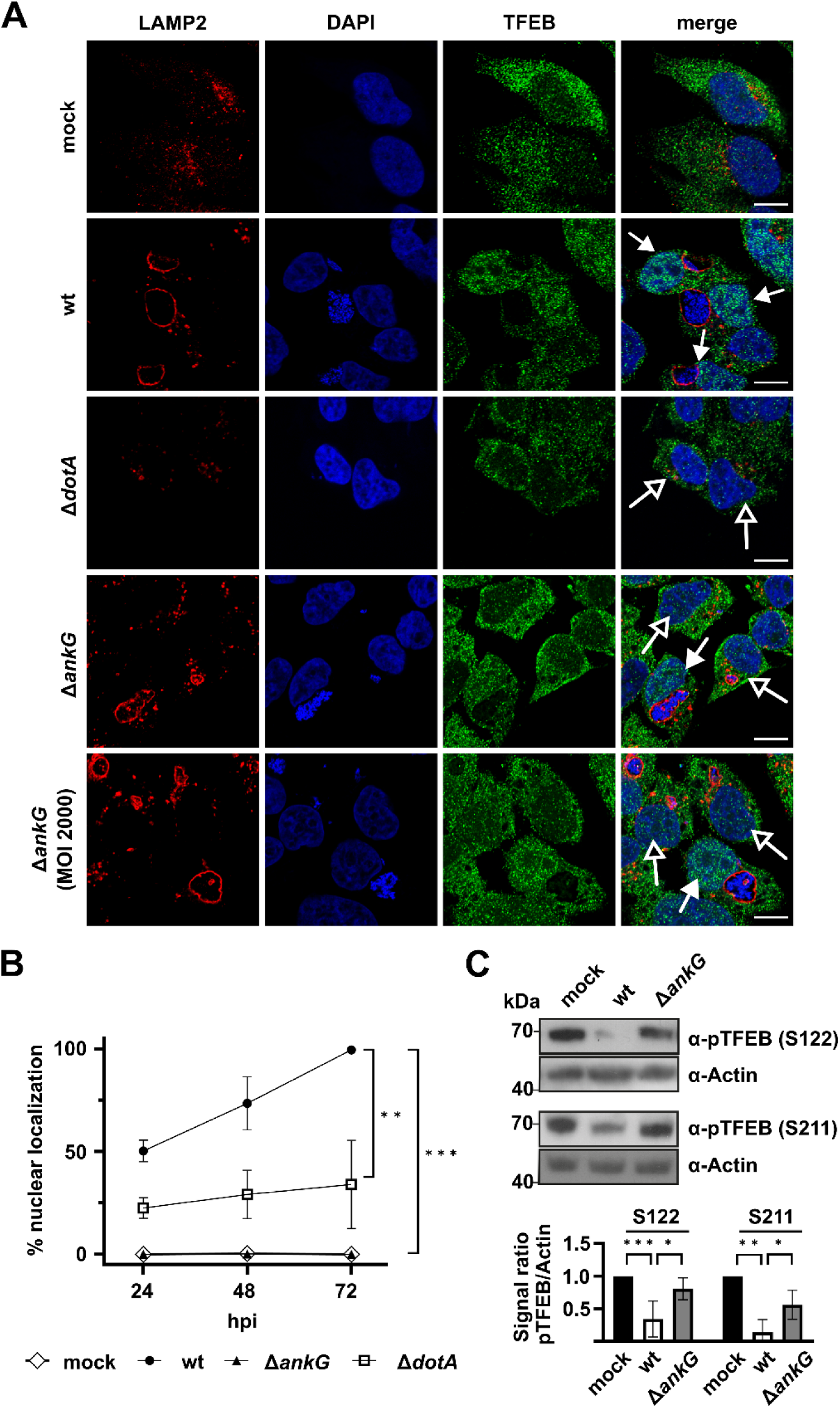
TFEB is activated by wild-type *C. burnetii* in a time-dependent manner. (A) HeLa cells were either not infected (mock) or infected with *C. burnetii* wild-type (wt), the T4BSS mutant Δ*dotA* or the Δ*ankG* mutant at MOI 200 for 72 hours. Cells were additionally infected with the Δ*ankG* mutant at MOI 2000. The cells were fixed, permeabilized and stained with antibodies against LAMP2 (red) and TFEB (green) and with DAPI (blue). The subcellular localization of TFEB was analyzed by confocal microscopy. Representative immunofluorescence images are shown. Open arrows indicate cytoplasmic localization of TFEB, while filled arrows indicate nuclear localization of TFEB. Scale bar = 10 µm. (B) HeLa cells were either not infected (mock) or infected with *C. burnetii* wild-type (wt), Δ*dotA* or Δ*ankG* at MOI 200 for 24, 48 and 72 hours. The cells were fixed, permeabilized and stained with an anti-TFEB antibody. One hundred cells were scored for nuclear localization of TFEB in each of three independent experiments. Error bars indicate ± SD. n=3, Mann-Whitney test. *** p < 0.001. (C) HeLa cells were either not infected (mock) or infected with *C. burnetii* wild-type (wt) or the Δ*ankG* mutant at MOI 200. 72 hours post-infection the cells were analyzed by Western blot analysis using antibodies against pTFEB (S122), pTFEB (S211) and Actin as loading control. Representative immunoblots out of three independent experiments with similar results are depicted. Densitometric analysis of the pTFEB/Actin ratio was performed using ImageJ. Arbitrary units (AU) are shown relative to uninfected cells (mock). Mean ± SD, n=3, one sample t-test and unpaired students t-test. *p < 0.05.

Activation of TFEB depends on nuclear localization, which in turn is controlled by its phosphorylation status. Phosphorylated TFEB is inactive as it interacts with 14-3-3 proteins, which results in cytoplasmic retention of TFEB (Martina *et al*., 2012). Especially phosphorylation at amino acids S122, S134, S142, S211, and S467 results in cytoplasmic retention (Puertollano *et al*., 2018). Here, the phosphorylation status of S122 and S211 in cells either not infected or infected with wild-type or the Δ*ankG* mutant was analyzed. While cells infected with the wild-type showed a clear reduced phosphorylation level at S122 and S211, cells infected with the Δ*ankG* mutant did not (Fig. 1C). These data demonstrate that *C. burnetii* induces dephosphorylation, and, thus, activation of TFEB. In contrast, the Δ*ankG* mutant is clearly defective in TFEB activation. There are two possible explanations for this outcome: i) Either the T4BSS effector protein AnkG is involved in TFEB activation or ii) the lack of the formation of a spacious CCV might prevent efficient activation of TFEB.

### AnkG does not directly activate TFEB

To test whether AnkG influences activation of TFEB, we ectopically expressed GFP-tagged AnkG and GFP as a negative control. However, the transfection *per se* resulted in nuclear migration of TFEB (data not shown). Therefore, a different approach was used. Thus, we established a HeLa Tet-On cell line inducibly expressing AnkG containing a nuclear localization signal (NLS). NLS-AnkG was used, as ectopically expressed AnkG has two different subcellular localizations, nuclear and in association with mitochondria, while NLS-AnkG is only localized within the host cell nucleus (Eckart *et al*., 2014). AnkG has to be within the host cell nucleus to fulfill its functional activity (Cordsmeier *et al*., 2022). In HeLa-pWHE644/655-NLS-AnkG cells, NLS-AnkG is expressed after addition of doxycycline (Fig. 2A). NLS-AnkG showed nuclear localization, as demonstrated by immunofluorescence staining (Fig. 2B). However, the expression of NLS-AnkG did not induce nuclear localization of TFEB (Fig. 2B), nor influence its phosphorylation status at S211 (Fig. 2A). In contrast, treatment with Torin 1 resulted in dephosphorylation of TFEB, confirming previous results (Napolitano et al., 2018). This dephosphorylation of TFEB (S211) also occured in HeLa-pWHE644/655-NLS-AnkG cells stimulated with doxycycline, demonstrating that the activation of TFEB in these cells was possible. Hence, AnkG is unable to induce dephosphorylation, and, thus, activation of TFEB. Consequently, we hypothesized that the reason for the reduced activation of TFEB in cells infected with the Δ*ankG* mutant might be due to altered vacuole size or form.

**Figure 2:**
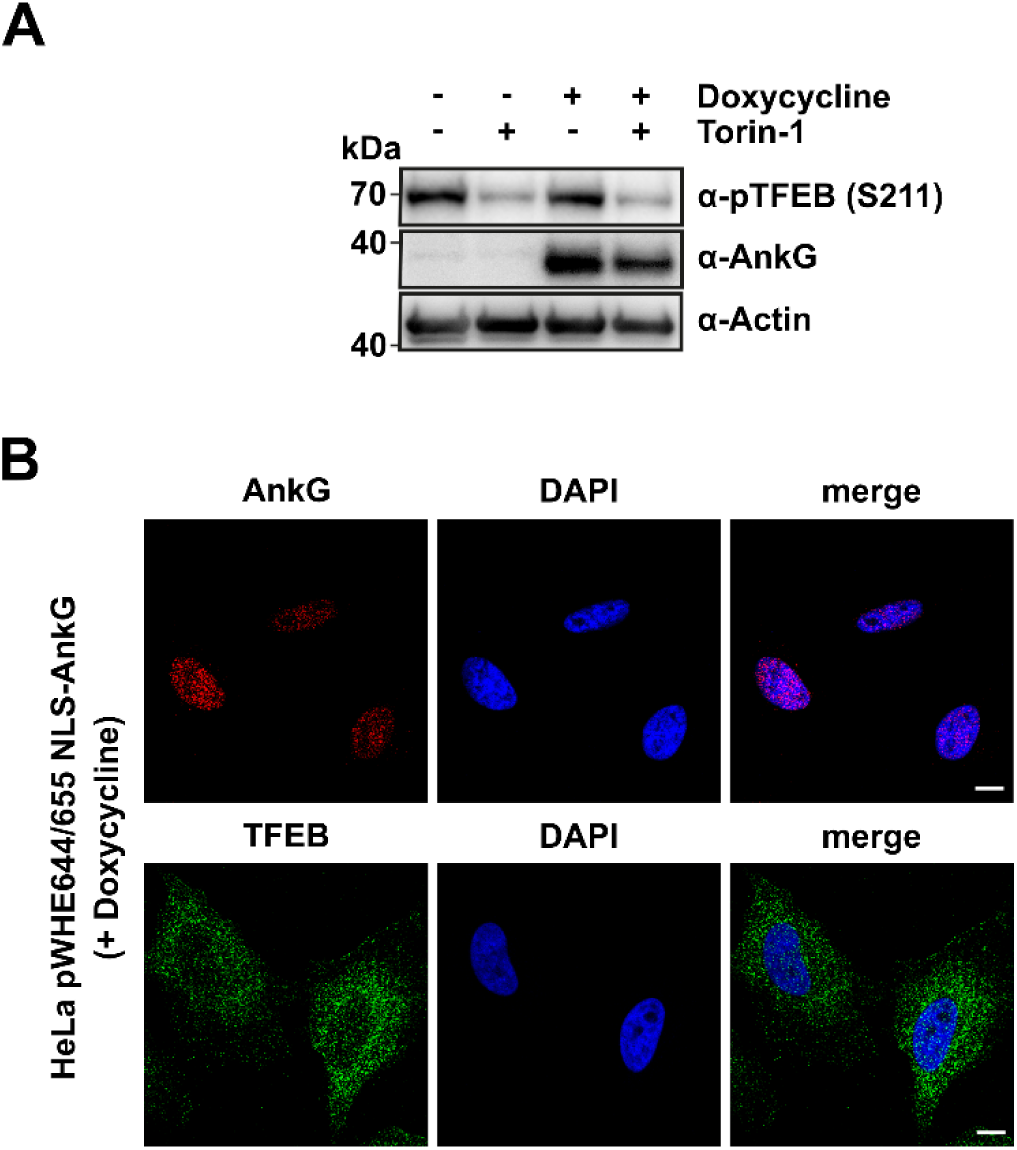
Ectopically expressed NLS-AnkG does not activate TFEB. HeLa cells were stably transfected with doxycycline inducible NLS-AnkG. (A) Cell lysates were analyzed by Western blot analysis using antibodies against AnkG, pTFEB (S211) and Actin as loading control. Representative immunoblots out of three independent experiments with similar results are shown. (B) The cells were fixed, permeabilized and stained with anti-TFEB and anti-AnkG antibodies and with DAPI. Representative immunofluorescence images are shown. Scale bar = 10 µm.

### Activation of TFEB correlates with the volume of the CCV

To challenge the above-mentioned hypothesis, the volume (in µm^3^) of the CCVs harboring wild-type or Δ*ankG C. burnetii* was determined at 72 hours post-infection. A significant difference between the sizes of Δ*ankG* and wild-type *C. burnetii-*containing vacuoles was observed. The wild-type formed a ∼5-fold larger CCV than the mutant (Fig. 3A). This suggests that only larger CCVs might induce TFEB activation. To verify this assumption, the CCV size of Δ*ankG C. burnetii* at 72 hours post-infection was correlated with the localization of TFEB. At this time point, only ∼30% of the cells infected with the Δ*ankG* mutant had nuclear, and, thus, activated TFEB (Fig. 1B). Indeed, the cells infected with the Δ*ankG* mutant which had nuclear TFEB also had larger CCVs (Fig. 3B). Similarly, in cells infected with wild-type *C. burnetii* for 24 hours, a time point at which 50% of the infected cells contained nuclear TFEB, the activation status of TFEB clearly correlated with larger CCV size (Fig. 3C). To further prove our hypothesis, the infection rate of the Δ*ankG* mutant was increased 10-fold. This resulted in increased CCV sizes (Fig. 3D) and concomitant dephosphorylation of TFEB (S211) (Fig. 3E), indicating activation of TFEB. From this data we concluded that the size of the CCV determines the activation of TFEB.

**Figure 3:**
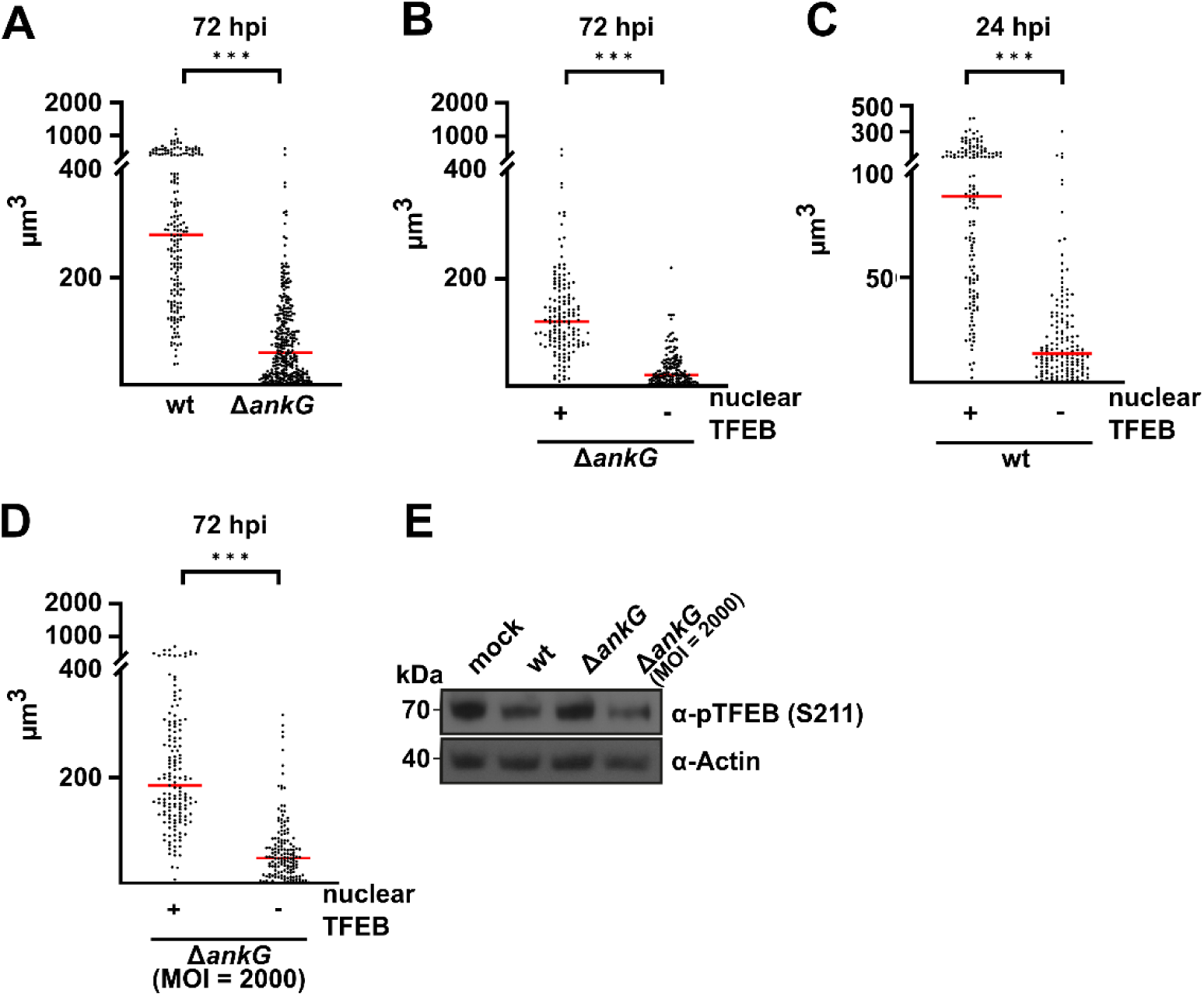
TFEB activation status correlates with the CCV size. HeLa cells were infected with wild-type *C. burnetii* (wt) or the Δ*ankG* mutant at MOI 200 for 24 and 72 hours. (A) The CCV sizes of cells infected with wild-type *C. burnetii* (wt) or the Δ*ankG* mutant for 72 hours were measured in combined confocal Z-stacks. The XYZ dimensions were analyzed with the profile module of the confocal software Zen 3.0 SR (black) (Zeiss). The volumes of at least 50 CCVs from each of three independent experiments are shown. The bar represents the mean. The Mann-Whitney test was performed with GraphPad Prism. *** p < 0.001. (B) The CCV sizes of cells infected with the Δ*ankG* mutant for 72 hours were measured and correlated with nuclear localization of TFEB. The volumes of at least 50 CCVs from each of three independent experiments are shown. The bar represents the mean. The Mann-Whitney test was performed with GraphPad Prism. *** p < 0.001. (C) The CCV sizes of cells infected with wild-type (wt) *C. burnetii* for 24 hours were measured and correlated with nuclear localization of TFEB. The volumes of at least 50 CCVs from each of three independent experiments are shown. The bar represents the mean. The Mann-Whitney test was performed with GraphPad Prism. *** p < 0.001. (D) HeLa cells were infected with the Δ*ankG* mutant at MOI 2000 for 72 hours. CCV sizes were measured and correlated with nuclear localization of TFEB. The volumes of at least 50 CCVs from each of three independent experiments are shown. The bar represents the mean. The Mann-Whitney test was performed with GraphPad Prism. *** p < 0.001. (E) HeLa cells were either not infected (mock) or infected with wild-type *C. burnetii* (wt) at MOI 200 and with the Δ*ankG* mutant at MOI 200 and 2000. 72 hours post-infection, the cells were analyzed by Western blot using an antibody against pTFEB (S211) and Actin as loading control. Representative immunoblots out of two independent experiments with similar results are depicted.

### Activation of TFEB in *C. burnetii* infected cells is mTORC1-independent

In the next step, we asked how TFEB is activated during infection. TFEB activity is primarily controlled by its subcellular localization, which is, in turn, regulated by post-translational modifications, such as phosphorylation. The phosphorylation of several serine residues keeps the protein in the cytosol. The main negative regulator is mTORC1, which phosphorylates S122, S142 and S211 in TFEB. However, other kinases are also known to phosphorylate TFEB, including GSK3b or PKB. It has been shown that inactivation of mTORC1 induces TFEB activation (Franco-Juarez *et al*., 2022). Importantly, it was shown that *C. burnetii* inhibits mTORC1 (Larson *et al*., 2019). For this reason, we analyzed the activation state of mTORC1 in our infection model. Hence, the phosphorylation patterns of mTOR, Raptor, PRAS40 and 4E-BP1 were analyzed, as downregulation of mTOR S2448 phosphorylation correlates with decreased mTORC1 activity (Rosner *et al*., 2010), phosphorylation at S792 in Raptor inhibits mTORC1 (Gwinn *et al*., 2010), phosphorylation of PRAS40 prevents its inhibition of mTORC1 (Sancak *et al*., 2007) and mTORC1 phosphorylates 4E-BP1 to stimulate translation (Ma *et al*., 2009). Dephosphorylation of mTOR (S2448) during infection was not detected (Fig. 4A), suggesting that mTOR is not inhibited during infection with wild-type *C. burnetii* or the Δ*ankG* mutant at the time periods analyzed. This is supported by the lack of increased phosphorylation of Raptor and increased phosphorylation of RPAS40 and 4E-BP1 during infection (Fig. 4A). We also analyzed the subcellular localization of mTORC1. Remarkably, mTORC1 activation takes place at the lysosomal membrane. Once mTORC1 is released from the lysosomal membrane it becomes inactive (Sancak *et al*., 2010). In *C. burnetii* infected cells we detected mTOR at the CCV membrane in ∼75% of the infected cells (Fig. 4B). As the CCV membrane resembles the lysosomal membrane, we concluded that mTORC1 is active in the infected cells. Taken together, the data suggest that mTORC1 is active, which suggest that TFEB activation in *C. burnetii*-infected cells is not mediated by inhibition of mTORC1.

**Figure 4:**
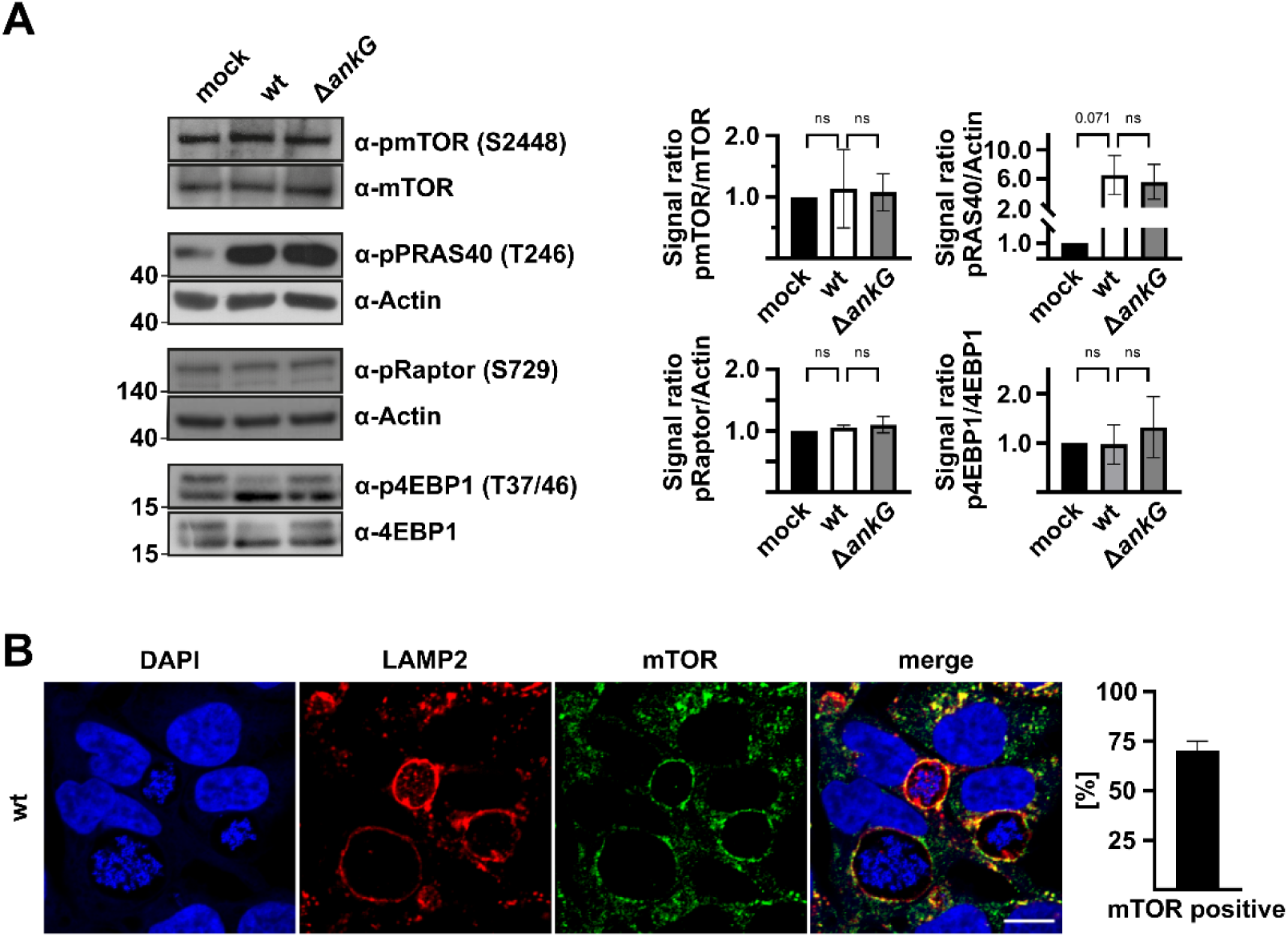
Infection with *C. burnetii* does not inhibit mTORC1 activation. (A) HeLa cells were either not infected (mock) or infected with *C. burnetii* wild-type (wt) or the Δ*ankG* mutant at MOI 200. 72 hours post-infection the cells were analyzed by Western blot using antibodies against pmTOR (S2448), pRaptor (S792), p4E-BP1 (T37/46), pPRAS40 (T246), and Actin, 4EBP1 or mTOR as loading controls. Representative immunoblots out of three independent experiments with similar results are shown. Densitometric analysis of the phosphorylated protein to loading control was performed using ImageJ. Arbitrary units (AU) are shown relative to uninfected cells (mock). Mean ± SD, n=3, one sample t-test and unpaired students t-test. ns= not significant, *p < 0.05. (B) HeLa cells infected with *C. burnetii* wild-type (wt) for 72 hours were fixed, permeabilized and stained with anti-mTOR and anti-LAMP2. Representative immunofluorescence images are shown. Scale bar = 10 µm. At least 50 CCVs in each of three independent experiments were analyzed for the co-localization with mTOR.

### MCOLN1 mediates activation of TFEB and CCV expansion

Activation of TFEB is mainly controlled by two phosphatases, Calcineurin and protein phosphatase 2A (PP2A). While Calcineurin is activated by Ca^2+^ or by calpain-mediated proteolysis (Klee *et al*., 1998, Tallant *et al*., 1988), PP2A is regulated at the level of expression, localization, holoenzyme composition and post-translational modification (Zolnierowicz, 2000). Consequently, alteration of Calcineurin or PP2A during *C. burnetii* infection was analyzed by immunoblot analysis. No influence of the infection on both phosphatases was detected at the protein level (Fig. 5A). This indicates that PP2A is not upregulated at the protein level, nor is Calcineurin cleaved during *C. burnetii* infection. Importantly, Calcineurin is also activated by release of lysosomal Ca^2+^ through the Ca^2+^-channel mucolipin 1 (MCOLN1) (Medina *et al*., 2015). Therefore, siRNA-mediated knock-down of MCOLN1 was performed, and the phosphorylation status of TFEB (S211) in cells either not infected or infected with wild-type *C. burnetii* was determined. As shown in figure 5B, the infection only resulted in reduced phosphorylation of TFEB (S211) in wild-type infected cells treated with non-targeting siRNA, but not in cells treated with MCOLN1 siRNA. In addition, reduced nuclear translocation of TFEB during infection in cells treated with MCOLN1 siRNA was observed (Fig. 5C). Hence, the presence of MCOLN1 is important for TFEB activation during *C. burnetii* infection. Consequently, we hypothesized that MCOLN1-mediated Ca^2+^ release leads to activation of Calcineurin and in turn to activation of TFEB, which is in line with previous results (Medina *et al*., 2015).

**Figure 5:**
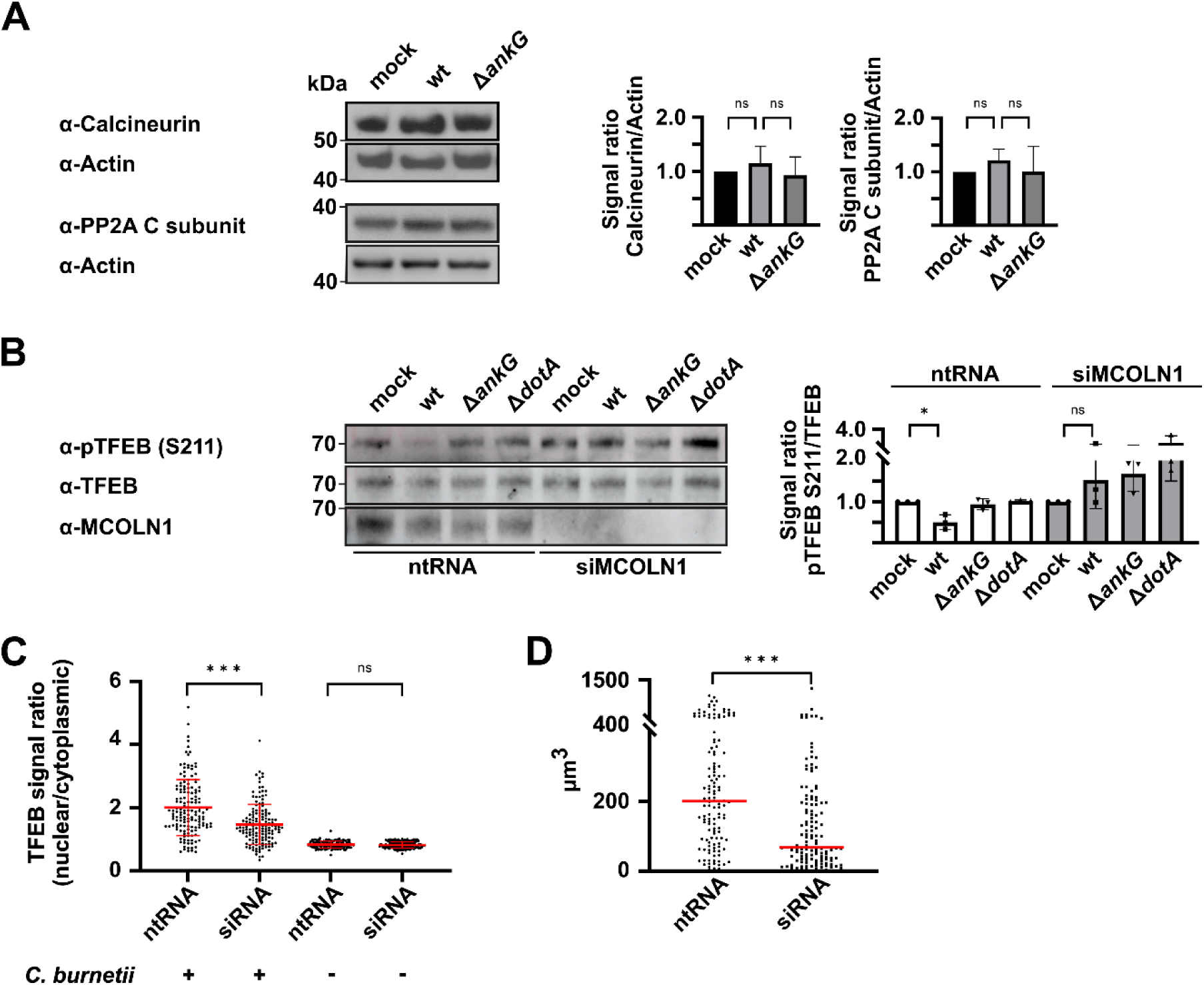
MCOLN1 and Calcineurin are important for TFEB activation during *C. burnetii* infection. (A) HeLa cells were either not infected (mock) or infected with wild-type *C. burnetii* (wt) or the Δ*ankG* mutant at MOI 200. 72 hours post-infection, the cell lysates were analyzed by Western blot using antibodies against Calcineurin, PP2A C subunit, and Actin as loading control. Representative immunoblots out of three independent experiments with similar results are depicted. Densitometric analysis of Calcineurin/Actin and PP2A C subunit/Actin ratios was performed using ImageJ. Arbitrary units (AU) are shown relative to uninfected cells (mock). Mean ± SD, n=3, one sample t-test and unpaired students t-test. ns= not significant. (B) HeLa cells were treated with ntRNA or siRNA targeting MCOLN1. 24 hours post-transfection, the cells were either not infected (mock) or infected with wild-type *C. burnetii* (wt), the Δ*ankG* or Δ*dotA* mutants at MOI 200. 48 hours post-infection the cell lysates were analyzed by Western blot using antibodies against MCOLN1, pTFEB (S211), and TFEB as loading control. Representative immunoblots out of three independent experiments with similar results are depicted. Densitometric analysis of pTFEB (S211)/TFEB ratio was performed using ImageJ. Arbitrary units (AU) are shown relative to uninfected cells (mock). Mean ± SD, n=3, one sample t-test. ns = not significant, ** p < 0.01. (C) HeLa cells, treated with ntRNA or siRNA targeting MCOLN1, were either infected or not with *C. burnetii* wild-type. Cells were fixed, permeabilized and labeled with anti-TFEB, anti-LAMP2 and DAPI. The ratio of nuclear and cytoplasmic TFEB was determined by comparing the pixel intensities of the different compartments using confocal microscopy and ImageJ. The ratio of nuclear to cytoplasmic TFEB was calculated from 50 cells in each of three independent experiments. The bar represents the mean. The Mann-Whitney test was performed with GraphPad Prism. ns = not significant, *** p < 0.001. (D) HeLa cells were treated with ntRNA or siRNA targeting MCOLN1. The cells were infected with *C. burnetii* at MOI 200. 48 hours post-infection, CCV sizes were measured. The volumes of at least 40 CCVs from each of three independent experiments are shown. The bar represents the mean. The Mann-Whitney test was performed with GraphPad Prism. *** p < 0.001.

To learn about the functional consequence of TFEB activation for CCV development, the size of the CCV harboring wild-type *C. burnetii* in cells treated with non-targeting siRNA or MCOLN1 siRNA was determined. A significant reduction in CCV size was observed when MCOLN1 was knocked-down. Thus, the CCV size not only induces TFEB activation (Fig. 3C), but MCOLN1-mediated activity is also required for CCV expansion (Fig. 5D).

### MCOLN1 activation results in egress of *C. burnetii*

It was previously shown that Galectin-3 is found on the CCV, possibly as a sensor of CCV membrane damage (Mansilla Pareja *et al*., 2017). The damaged CCV membrane seems to be repaired by reversible recruitment of the ESCRT machinery (Radulovic *et al*., 2018), for which ALIX is important (McCullough *et al*., 2018). However, the *C. burnetii* T4BSS effector protein Vice prevents efficient recruitment of ALIX to the CCV (Bienvenu *et al*., 2024). Therefore, we hypothesized that CCV expansion might cause CCV membrane damage, which might not be efficiently repaired, and opens the possibility that Ca^2+^ leaks out of the injured CCVs. To get a first impression whether CCV membrane leakage might occur, the protein level of Galectin-3 during *C. burnetii* infection was analyzed. As shown in figure 6A, the protein level of Galectin-3 was upregulated by infection with *C. burnetii*, which indicates that the cell possibly reacts to membrane damage. Thus, it might be possible that with increasing size of the CCV, the membrane integrity might become constrained, which results in increased release of Ca^2+^ into the host cell cytosol. Elevated cytosolic Ca^2+^ levels either mediated by lysosomal membrane damage or by release via the MCOLN1 channel, might facilitate the fusion of lysosomes with the plasma membrane (Tardieux *et al*., 1992, Tancini *et al*., 2020). Therefore, we hypothesized that interference with MCOLN1-mediated Ca^2+^ release might alter lysosomal exocytosis. To test this hypothesis, HeLa cells infected for 24 hours, a time point, at which *C. burnetii* egress was not observed (Schulze-Luehrmann *et al*., 2024), were treated with either ML-SA1 to activate MCOLN1 (Shen *et al*., 2012) or with Torin 1 to activate TFEB by inhibiting mTOR (Thoreen *et al*., 2009) or left untreated. While no difference in the intracellular bacterial load was observed between the differently treated cells, an increased bacterial release into the supernatant was detected following ML-SA1 treatment (Fig. 6B). Importantly, treatment with Torin 1, which leads to TFEB activation, did not lead to augmented release of bacteria. This suggests, that ML-SA1-mediated activation of MCOLN1, and, thus, increased Ca^2+^ release from lysosomes results in *C. burnetii* egress.

**Figure 6:**
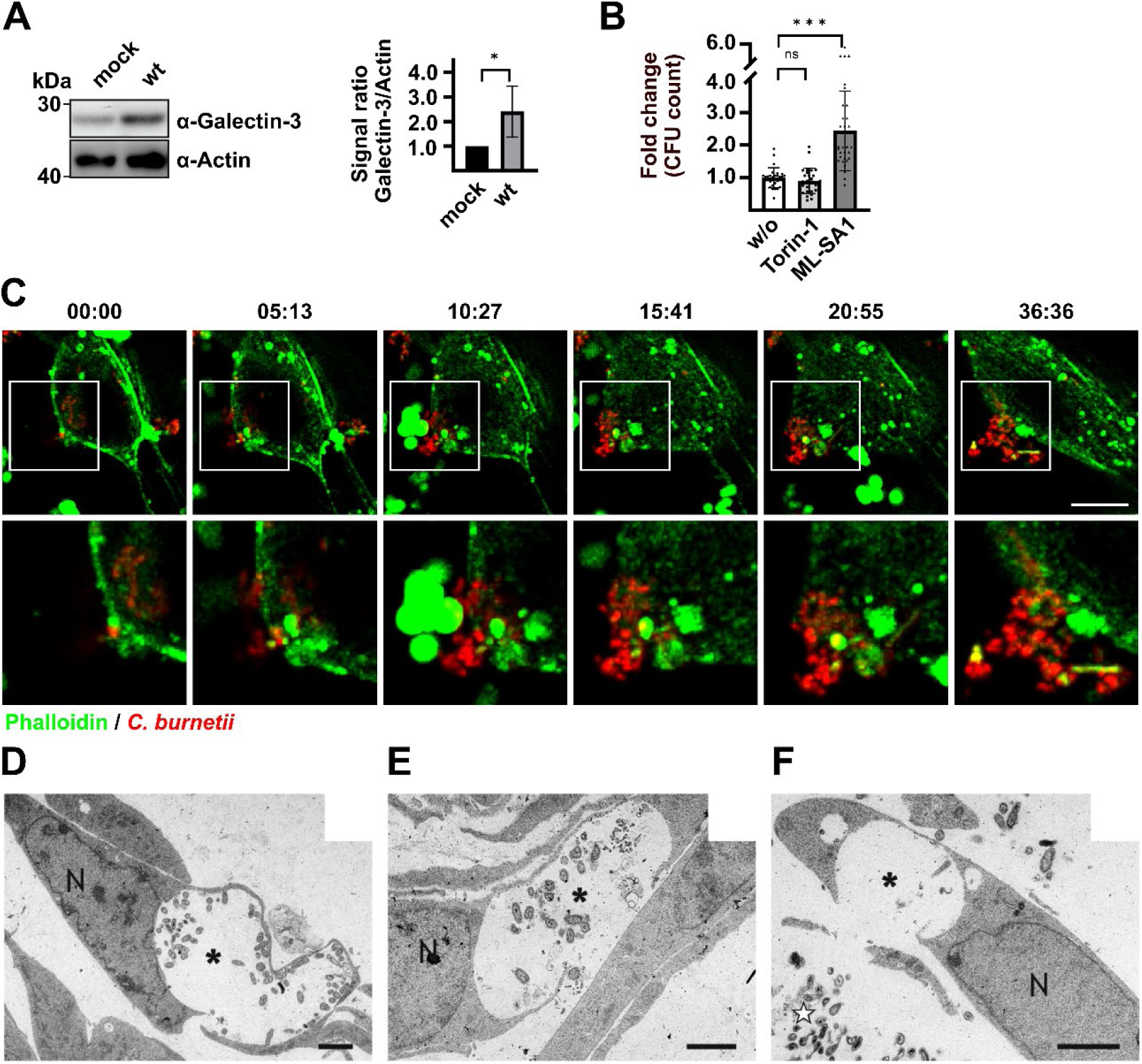
*C. burnetii* egresses via lysosomal exocytosis. (A) HeLa cells were either not infected (mock) or infected with wild-type *C. burnetii* (wt) at MOI 200. Cell lysates were analyzed by Western blot using antibodies against Galectin-3 and Actin, the latter serving as loading control. Representative immunoblots out of three independent experiments with similar results are depicted. Densitometric analysis of the Galectin-3/Actin ratio was performed using ImageJ. Arbitrary units (AU) are shown relative to uninfected cells (mock). Mean ± SD, n=3, one sample t-test and unpaired students t-test. ** p < 0.01. (B) HeLa cells were infected with wild-type *C. burnetii* at MOI 200 for 6 h. Cells were washed ten times and were treated without (w/o), with Torin 1 (250 nM) or with ML-SA1 (20 µM) for 18 hours. The CFU counts of the supernatants from three independent experiments performed in triplicates are shown. The fold change in CFU relative to untreated cells (w/o) are shown. The bar represents the mean. The Mann-Whitney test was performed with GraphPad Prism. ns= not significant, *** p < 0.001. (C) EA.hy926 cells were infected with *C. burnetii* at MOI of 200. After 24 h of infection, the cells were washed five times with PBS and treated with gentamicin (200 µg/ml) for 4 h to eliminate extracellular bacteria. Five days post-infection, the cells were visualized by spinning disc microscopy. The actin cytoskeleton was stained with SiR-actin (green). Images were acquired every ∼5 min. Time stamp: min:sec. Scale bar: 10 µm. The white inlet shows a 4-fold magnification. (D-F) Wild-type MEFs were infected for 7 days with *C. burnetii*. The cells were fixed with glutaraldehyde, embedded in araldite and ultrathin sectioned for transmission electron microscopy. Open CCVs (marked with *) release *C. burnetii* into the extracellular space (marked with white stars). All cells affected have normal nuclear (marked with N) morphology and no signs of cellular damage. The opening of the CCV in (D) is small, larger in (E) and extensive in (F). While in (D) and (E), *C. burnetii* are still found in the CCVs, the CCV in (F) is empty, with numerous bacteria in the extracellular space. Scale bar: 2 µm.

### *C. burnetii* egresses via lysosomal exocytosis

We have previously shown that apoptosis-induction at later stages of infection allows *C. burnetii* egress (Schulze-Luehrmann *et al*., 2024). Lysosomal exocytosis leads to plasma membrane repair, and, thus, survival of the cell (Reddy *et al*., 2001). To follow the hypothesis that Ca^2+^ release from lysosomes and/or CCVs either via the Ca^2+^ channel MCOLN1 or by membrane damage triggers lysosomal exocytosis, live cell imaging at later stages of infection was performed. Indeed, at 5 days post-infection infected cells were observed, in which *C. burnetii* leaks out of the cells without obvious cell death induction (Fig. 6C). This indicates that *C. burnetii* might also use a non-apoptotic pathway to leave its host cell. In agreement with this hypothesis, release of *C. burnetii* from mouse embryonic fibroblasts (MEFs) with healthy nuclei into the extracellular space was observed after 7 days of infection (Fig. 6D-E). Together, this opens the possibility that lysosomal exocytosis might be a possible egress pathway.

### LAMP1/2 is essential for lysosomal exocytosis-mediated egress of *C. burnetii*

Lysosomal exocytosis is controlled by the subcellular localization of lysosomes, as lysosomes have to move towards the plasma membrane (Tancini *et al*., 2020). Peripheral lysosomes are more alkaline than juxtanuclear/central lysosomes (Sbano *et al*., 2017, Johnson *et al*., 2016). Consequently, the pH values of CCVs were measured and correlated with their subcellular localization, juxtanuclear/central versus peripheral. Indeed, the pH value of peripheral CCVs was more alkaline than that of juxtanuclear/central CCVs (Fig. 7A), opening the possibility that the peripheral CCVs might undergo lysosomal exocytosis.

**Figure 7:**
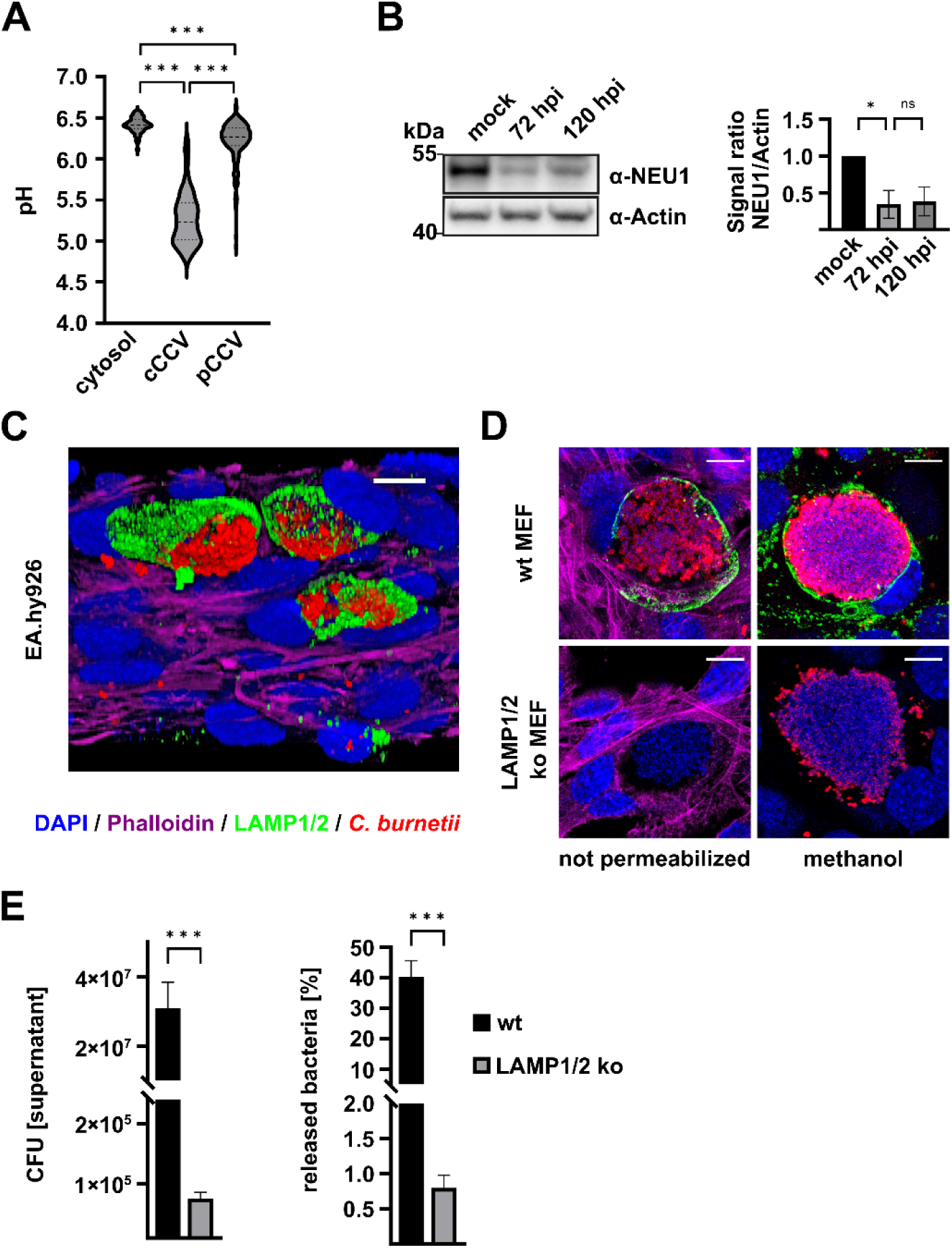
LAMP1/2 are required for lysosomal exocytosis of *C. burnetii*. (A) Quantification of cytosolic and vesicular pH (see Experimental Procedures) of the perinuclear/central (cCCV) and peripheral (pCCV) CCVs 5d post-infection in EA.hy926 cells based on fluorescence intensity ratios of the dual-wavelength fluorophore LysoSensor Yellow/Blue DND-160. Data represent average values ± SD of 30 infected cells per sample in duplicates from three independent experiments. Ordinary one-way ANOVA with Tukeýs multiple comparisons were performed with GraphPad Prism. *** p < 0.001. (B) HeLa cells were either not infected (mock) or infected with *C. burnetii* at MOI 200 for 72 hours (72 hpi) or 120 hours (120 hpi). Cell lysates were analyzed by Western blot using antibodies against NEU1 and Actin as loading control. Representative immunoblots out of three independent experiments with similar results are depicted. Densitometric analysis of the NEU1/Actin ratio was performed using ImageJ. Arbitrary units (AU) are shown relative to uninfected cells (mock). The bar represents the mean. One sample t-test and unpaired students t-test was performed with GraphPad Prism. ns= not significant, ** p < 0.01. (C) EA.hy926 cells were infected with *C. burnetii* at MOI 200. After 24 h of infection, the cells were washed five times with PBS and treated with gentamicin (200 µg/ml) for 4 h to eliminate extracellular bacteria. Five days post-infection, cells were visualized by confocal microscopy without permeabilization. The cells were stained with Phalloidin (to mark actin; purple), anti-LAMP (green) and anti-*Coxiella* (red). (D and E) Wild-type MEFs (WT) and LAMP1/2 knock-out MEFs were infected with *C. burnetii* for 96 hours. (D) The cells were either permeabilized with Methanol or not and stained with Phalloidin (purple), anti-LAMP (green), anti-*Coxiella* (red) and DAPI (blue). Scale bar: 10 µm. (E) The CFU counts of the supernatant and percentage of released bacteria out of the total number of intracellular bacteria and in the supernatant from two independent experiments performed in triplicate are shown. The bar represents the mean. A paired t-test was performed with GraphPad Prism. *** p < 0.001.

Neuraminidase-1 (NEU1) is a sialidase, that modifies many membrane receptors to either activate or inhibit their function. Importantly, the post-transcriptional modulation of LAMP1 by NEU1 impairs lysosomal exocytosis (Yogalingam *et al*., 2008). The infection with *C. burnetii* resulted in reduced protein levels of NEU1 (Fig. 7B), which might abrogate the inhibition of lysosomal exocytosis. Thus, the data presented so far supports the possibility that *C. burnetii* uses lysosomal exocytosis as a pathway to exit its host cells.

To further verify that *C. burnetii* egress via lysosomal exocytosis, immunofluorescence microscopy of EA.hy926 cells infected with *C. burnetii* for 5 days without permeabilization was performed (Andrews, 2017). Importantly, the antibodies will only stain the bacteria if the CCV has access to the environment, which would be the case if the CCV is in the process of lysosomal exocytosis. Indeed, the bacteria in several infected cells were stainable. In addition, LAMP1/2 was also stained with an antibody in these cells (Fig. 7C). As the antibody binds to a luminal part of LAMP1/2, it indicates that the CCV has access to the environment. Together, these data demonstrate that *C. burnetii* egresses via lysosomal exocytosis.

There are several reports which suggest that LAMP2 plays an important role in lysosomal exocytosis (Couto *et al*., 2017, Grochowska *et al*., 2023, Kima *et al*., 2000, Yogalingam *et al*., 2008). Thus, the ability of *C. burnetii* to egress from LAMP1/2 knock-out MEFs and corresponding wild-type MEFs was determined (Eskelinen *et al*., 2004). Importantly, *C. burnetii* invade MEFs independent of LAMP1/2. Bacterial load is ∼ 2-fold higher in LAMP1/2 ko MEFs than in wild-type MEFs (Schulze-Luehrmann *et al*., 2016). At 4 days post-infection, *C. burnetii* was stainable with an antibody in non-permeabilized wild-type MEFs, but not in LAMP1/2 ko MEFS (Fig. 7D), indicating that only CCVs in the wild-type MEFs might undergo exocytosis. In order to confirm this, a CFU assay from the supernatants of infected wild-type and LAMP1/2 ko MEFs were performed. A more than 2 log-decrease in viable bacteria was detected in the supernatant of LAMP1/2 ko MEFs (Fig. 7E), indicating that LAMP1/2 is essential for lysosomal exocytosis of *C. burnetii*.

## DISCUSSION

*C. burnetii* is so far the only known bacterial pathogen which replicates within a phagolysosomal-like compartment. Lysosomes are the endpoint of the phagocytic pathway, and thereby essential to clear a bacterial infection. However, *C. burnetii* even requires the acidification within the CCV for bacterial metabolism, replication and translocation of T4BSS effector proteins into the host cell cytosol (Omsland *et al*., 2009, Newton *et al*., 2013). Nevertheless, *C. burnetii* seems to be able to reduce the number of proteolytically active vesicles within the host cell (Samanta *et al*., 2019), suggesting that *C. burnetii* is able to balance the activity of lysosome function. One way to regulate lysosomal biogenesis is via the transcription factor TFEB. The information provided so far concerning the function of TFEB during *C. burnetii* infection is quite diverse (Killips *et al*., 2024, Larson *et al*., 2019, Padmanabhan *et al*., 2020), which might be caused by the different tools used, overexpression of GFP-tagged TFEB versus TFEB ko versus endogenous TFEB. Here, we investigated the subcellular localization and activation status of endogenous TFEB, as this might be less sensitive to unwanted side-effects. Activation of endogenous TFEB during *C. burnetii* infection was observed in a time-dependent manner (Fig. 1), which is determined by the size of the CCV. Cells containing a large CCV are most likely characterized by active TFEB (Fig. 3). It was discussed previously that TFEB activation depends on the T4BSS (Padmanabhan *et al*., 2020). Although, we did not detect activation of TFEB in cells infected with a T4BSS defective mutant and only reduced activation of TFEB in cells infected with the Δ*ankG* mutant (Fig. 1A-C), we concluded that this might be due to the nature of the CCV generated by these two mutants. The T4BSS mutant is unable to generate a large CCV (Beare *et al*., 2011, Carey *et al*., 2011) and the Δ*ankG* mutant establishes smaller CCVs (Cordsmeier *et al*., 2022). Indeed, ectopic expression of the T4BSS effector protein AnkG in the absence of infection did not activate TFEB (Fig. 2), indicating that AnkG only indirectly activates TFEB by influencing CCV size. Similarly, we speculate that the fact that the T4BSS mutant did not activate TFEB at all, is a consequence of the mutant’s inability to establish a growing CCV. Further experiments are needed in the future to dissect the role of the T4BSS in TFEB activation.

Lysosomes not only play a role as a degradative organelle containing a wide repertoire of acid hydrolases, they are also important signaling platforms. Multiple cellular stress responses are initiated at the lysosomal membrane (Saftig *et al*., 2021). For regulation of cellular metabolism mTORC1 is important. Under ample supply of amino acids and/ or glucose, the mTORC1 complex is recruited to the lysosomal membrane, while under starvation conditions, mTORC1 shuttles to the host cell cytosol (Saftig *et al*., 2021, Lawrence *et al*., 2019). It was recently shown that during amino acid starvation, the infection with *C. burnetii* triggers increased inhibition of mTORC1. This *C. burnetii*-mediated mTORC1-inhibtion is maintained when nutrients were replenished. Importantly, the *C. burnetii*-mediated inhibition of mTORC1 seems to depend on the T4BSS (Larson *et al*., 2019). Here, we did not detect an inhibition of mTORC1 during *C. burnetii* infection, which might be due to the fact that complete media with high glucose was used (Fig. 4). Together these data suggest, that in the present of sufficient nutrients, *C. burnetii* might not trigger mTORC1 inhibition. Once nutrients become lacking, not only will the host cell inhibit mTORC1 to adapt to this limitation, but also the pathogen. How *C. burnetii* sense a lack of nutrients is unknown, as well as the signaling cascade leading to mTORC1 inhibition, but T4BSS effector protein(s) might be involved (Larson *et al*., 2019).

TFEB is recruited to the lysosomal membrane under basal conditions (Martina *et al*., 2013). This allows phosphorylation, and, thus, inactivation by mTORC1. Under nutrient limiting conditions, mTORC1 is inhibited, which results in TFEB dephosphorylation and activation (Martina *et al*., 2014). However, our experiments showed that during *C. burnetii* infection under ample nutrient supply, activation of TFEB is not a result of mTORC1 inhibition (Fig. 4), which is in line with a previous report (Larson *et al*., 2019). Activation of TFEB by dephosphorylation is not only mediated by mTORC1 inactivation, but also by the activity of the phosphatases Calcineurin (Medina *et al*., 2015) and PP2A (Martina *et al*., 2018). Indeed, we could show that TFEB activation during *C. burnetii*-infection depends on the Ca^2+^ channel MCOLN1 (Fig. 5).

Here, we propose that with increasing CCV sizes not only is TFEB activated, but the CCV membrane might also be damaged. Thus, Galectin-3 protein levels were increased at later stages of a *C. burnetii* infection (Fig. 6A), which is an indication for lysosomal membrane damage. Damage of the lysosome or, in this case of the CCV, a phagolysosomal-like compartment, results in increased cytoplasmic Ca^2+^ levels, which might trigger bacterial egress via lysosomal exocytosis (Tancini *et al*., 2020, Tardieux *et al*., 1992). For the establishment of disease, it is crucial that intracellular pathogens are able to spread from their first target cells to other cells, cell types and organs. After completion of their replication cycle the bacteria have to egress, which is a complex process, and depend on the interplay between pathogen and host. There are several pathways described leading to egress, which either depend on induction of host cell death, extrusion, protrusion or exocytosis (Flieger *et al*., 2018). For the obligate intracellular pathogen *C. burnetii* knowledge about egress strategies is limited. We have previously shown that induction of apoptosis at later stages of infection is involved in egress and spreading of *C. burnetii*, but other pathways have to be employed, too (Schulze-Luehrmann *et al*., 2024). Here, we showed that *C. burnetii* egress from their host cell via lysosomal exocytosis (Fig. 6C-F and 7). Lysosomal exocytosis of *C. burnetii* correlates with a decreased NEU1 protein level (Fig. 7B), which leads to increased numbers of LAMP1-positive lysosomes at the plasma-membrane and extracellular release of lysosomal content (Yogalingam *et al*., 2008). In addition, NEU1 downregulation curtails pro-inflammatory responses (Lillehoj *et al*., 2022), suggesting that *C. burnetii*-mediated reduction of the NEU1 protein level might not only be important for bacterial exocytosis, but also for modulating immune responses. How *C. burnetii* influence the NEU1 protein level and how this affects the host reaction to the infection requires further investigations. Importantly, *C. burnetii* exocytosis depends on the presence of LAMP1/2 (Fig. 7). Although *C. burnetii* replicates in LAMP1/2 ko MEF to a higher number as in wild-type (Schulze-Luehrmann *et al*., 2016), we observed a strong reduction in released bacteria (Fig. 7E).

TFEB activation by ML-SA1-mediated activation of MCOLN1 resulted in lysosomal exocytosis, while Torin 1-mediated TFEB activation did not (Fig. 6B). This indicates that there is probably a different layer of control of TFEB activation, either mediated by the host and/ or the pathogen. Further research is required to understand how *C. burnetii* uses TFEB action for the own benefit and how interfering with this process might be useful to control the infection.

## ACKNOWLEDGEMENTS

We thank Dr. Alfonso Felipe-Lopez for image processing of figure 6C, Dr. Stephanie Bisle-Kitsiou for generating the HeLa-pWHE644/655-AnkG cell line, and Dr. Christian Berens for critical reading of the manuscript.

## AUTHOR CONTRIBUTIONS

AL - conceptualization; AL - funding acquisition; AL – supervision; SR, JSL, FW, ELT – investigation and analysis; AL, SR, JSL - writing first draft; all – writing and editing.

